# High contribution of *Pelagibacterales* to bacterial community composition and activity in spring blooms off Kerguelen Island (Southern Ocean)

**DOI:** 10.1101/633925

**Authors:** J. Dinasquet, M. Landa, I. Obernosterer

**Affiliations:** CNRS, Sorbonne Université, Laboratoire d’Océanographie Microbienne, LOMIC, F-66650 Banyuls-sur-Mer, France; Marine biology research division and Climate, Atmospheric Science & Physical Oceanography department, Scripps Institution of Oceanography, San Diego, USA; Ocean Sciences Department, University of California, Santa Cruz, USA

**Keywords:** iron fertilization, SAR11 clade, ecotypes, bacterial community composition, Kerguelen Plateau

## Abstract

The ecology of Pelagibacterales (SAR11 clade), the most abundant bacterial group in the ocean, has been intensively studied in temperate and tropical ocean regions, but the distribution patterns of this clade remains largely unexplored in the Southern Ocean. Through amplicon sequencing of the 16S rRNA genes, we assessed the contribution of Pelagibacterales to bacterial community composition in the naturally iron fertilized region off Kerguelen Island (Southern Ocean). We investigated the upper 300 m water column at seven sites located in early spring phytoplankton blooms and at one site in HNLC waters. Despite pronounced vertical patterns of the bacterioplankton assemblages, the SAR11 clade had high relative abundances at all depths and sites, averaging 40% (±15%) of the total community relative abundance. Micro-autoradiography combined with CARD-FISH further revealed that the SAR11 clade contributed substantially (45-60% in surface waters) to bacterial biomass production (as determined by ^3^H leucine incorporation). A clear niche partitioning of the further resolved SAR11 subclades was observed with depth layers, but differences among sites were detectable for only a few subclades. Our study provides novel observations of the distribution and contribution to the marine carbon cycle of the SAR11 clade in the cold waters of the Southern Ocean.

## Introduction

The SAR11 clade of the alphaproteobacteria is one of the most abundant bacterioplankton in marine ecosystems (Morris *et al.*, 2002; Carlson *et al.*, 2009; Eiler *et al.*, 2009) representing 25% or more of the total bacterial cells in seawater worldwide (Giebel *et al.*, 2009; Brown *et al.*, 2012; Sunagawa *et al.*, 2015; Ortmann and Santos, 2016). The reason of its success is still unclear (Brown *et al.*, 2014), but since its first discovery (Giovannoni *et al.*, 1990) a better understanding of its spatial, vertical and seasonal patterns and links to ecosystem variables has now emerged (e.g. Carlson *et al.*, 2009; Eiler *et al.*, 2009; Brown *et al.*, 2012; Morris *et al.*, 2012; Vergin *et al.*, 2013; Salter *et al.*, 2014; Thrash *et al.*, 2014; Ortmann and Santos, 2016). From these new insights into the clade’s ecology, several phylogenetic subclades with recurrent occurrence in specific environmental conditions have been identified (e.g. Field *et al.*, 1997; Carlson *et al.*, 2009; Vergin *et al.*, 2013). These subclades may represent ecologically coherent populations or ecotypes as suggested by their different traits and metabolic needs (Grote *et al.*, 2012). Hence, considering that SAR11 may be the most successful bacterioplankton lineage and thus, play a key role in marine biogeochemical cycles it is essential to provide new details on its population dynamics and subclades partitioning.

While members of the SAR11 clade have been found everywhere in the ocean, there are fewer studies on SAR11 subclades distribution and ecology in the Southern Ocean compared to temperate and tropical marine systems (Brown *et al*., 2012). The Southern Ocean presents a particular habitat for autotrophic and heterotrophic microbial communities, as the essential element iron (Fe) is present at low concentrations and therefore limits biological activities and the drawdown of carbon dioxide (CO_2_) in surface waters of High Nutrient Low Chlorophyll (HNLC) regions (Blain *et al.*, 2007; Pollard *et al.*, 2009). Further, concentrations of dissolved organic carbon (DOC) in Southern Ocean surface waters are among the lowest of the global ocean (about 50 µM; Hansell et al. 2010), representing an additional constraint for microbial heterotrophic activity. Whether Fe or organic carbon limit heterotrophic microbes varies on spatial and temporal scales in the Southern Ocean (summarized in Obernosterer *et al.*, 2015), and this likely reflects the tight links between autotrophs and heterotrophs through these key resources. On the one hand, the requirements of Fe for bacterial heterotrophic metabolism and growth (Tortell *et al.*, 1996; Fourquez *et al.*, 2014; Baltar *et al.*, 2018; Koedooder *et al.*, 2018) can lead to a potential competition with phytoplankton for this micronutrient (Fourquez *et al.*, 2016). On the other hand, the enhanced production of dissolved organic matter (DOM) by phytoplankton upon Fe-addition can stimulate bacterial metabolism. This interplay between the competition for a scarce resource and the response to phytoplankton-derived bioavailable DOM could influence the activity and abundances of bacterial taxa, including the SAR11 clade.

In the present study, we describe the spatial distribution of bacterial assemblages in the Kerguelen area of the Southern Ocean during a mosaic of phytoplankton blooms induced by natural Fe fertilization. The results show that the ubiquitous SAR11 clade was dominant in early spring, and actively contributing to bacterial carbon utilization both in HNLC waters and at the onset of phytoplankton blooms. We further resolved the SAR11 population dynamics to subclades showing niche partitioning with depth layers and bloom regimes.

## Results and discussion

The samples for the present study were collected in the naturally Fe-fertilized and in high nutrient low chlorophyll (HNLC) waters off Kerguelen Island in early spring during the KEOPS2 cruise (Fig.S1). The detailed hydrographic conditions are described in Park *et al.* (2014) and d’Ovidio *et al.* (2015). Concentrations of Chl *a*, bacterial abundance and bacterial heterotrophic production were overall higher in surface and intermediate waters of the naturally Fe-fertilized region as compared to the HNLC station R-2 (Christaki *et al.*, 2014; Lasbleiz *et al.*, 2014; Quéroué *et al.*, 2015) (TableS1). However, among the Fe-fertilized sites, considerable variability in these biological parameters were observed, reflecting spatial and temporal variability in the blooms development (Lasbleiz *et al.*, 2016) with Chl *a* concentrations ranging from 1.0 to 5.1 µg L^−1^ (Table S1). By contrast, the major inorganic nutrients N and P and DOC were similar across sites and characteristic for this region (Blain *et al.*, 2015; Tremblay *et al.*, 2015). In the present study, we focus on three water layers named surface, intermediate and deep (Table S1), identified by cluster analyses of abiotic and biotic environmental parameters (SIMPROF test difference between clusters of depth layers p<0.05, Fig. 1A).

**Figure 1:**
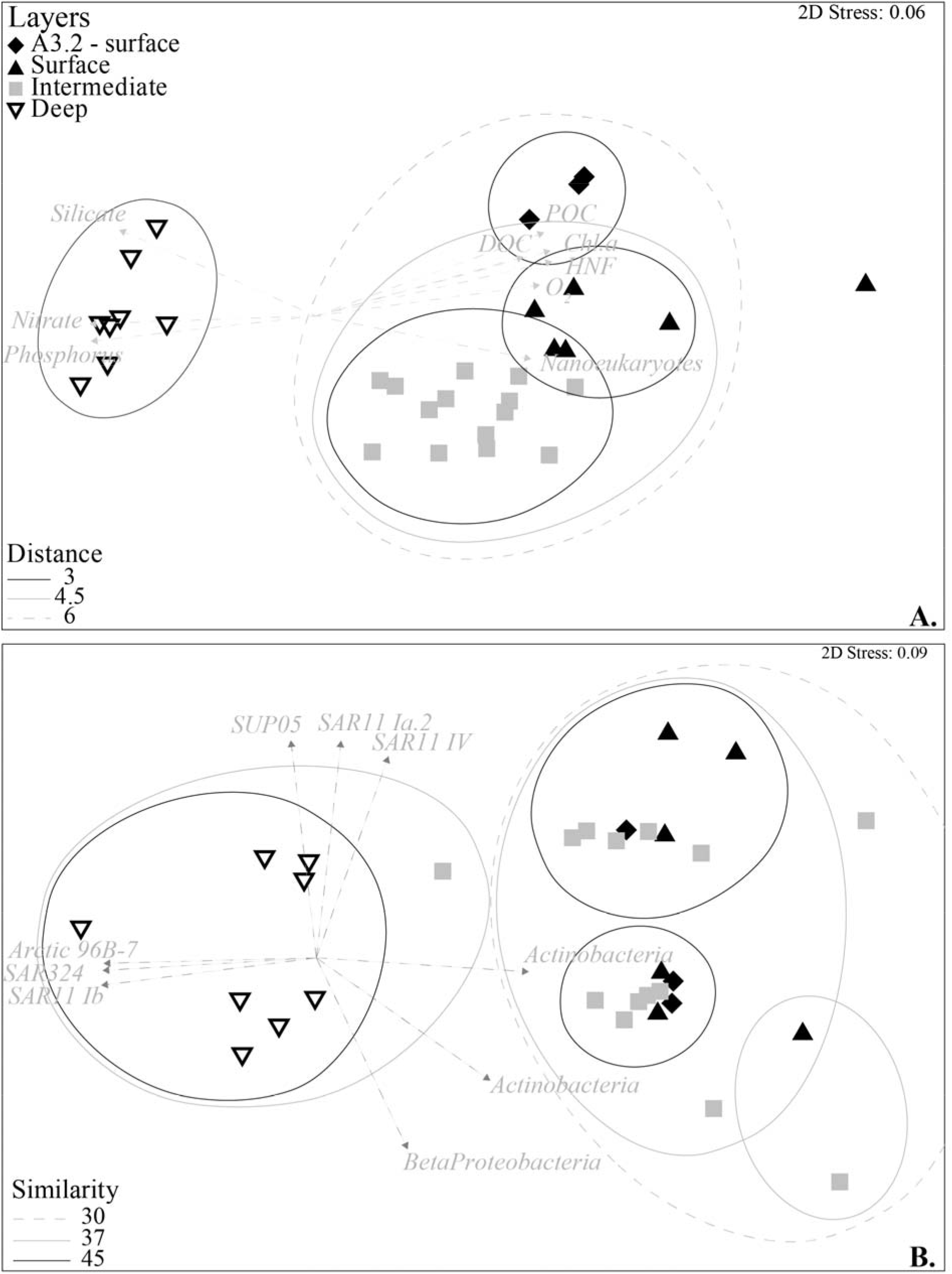
Depth layer ordination of the samples by environmental parameters (A), nMDS plot of all stations/depths organized by layers based on Euclidan distances of Log X+1 normalized environmental data. SIMPROF test showed significant differences between clusters (p<0.05) of samples representing deep, intermediate and surface layers. Vectors represent environmental parameters predicting nMDS axis with Spearman rank correlation > 0.85. Depth layer ordination of the samples by bacterial community composition (B), nMDS plot of all stations/depths organized by layers based on Bray-Curtis similarity index of log X+1 bacterial community dataset, vectors represent OTU (as closest relative taxonomic affiliation) predicting nMDS axis with Spearman rank correlation > 0.85. DOC – Dissolved Organic carbon; POC-Particulate Organic Carbon; O_2_-Dissolved oxygen concentration; HNF – Abundance of heterotrophic nanoflagellates.

The overall bacterial community composition based on the 16S rRNA gene, was significantly different (p<0.05) between the shallower and deep layers (Fig. 1B). However, no significant differences were observed between surface and intermediate layers. Richness and diversity were overall similar across sites in surface and intermediate layers, but these indices increased with depth at all stations (Table. S1). Proteobacteria were dominant in most of the stations/depth layers (on average 72 ±20% of total OTUs relative abundance), Actinobacteria were abundant in depth layers <150 m (up to 40%) and Bacteroidetes (mainly Flavobacteria) were relatively abundant in the surface layers (13 ±6%). Proteobacterial subclass distribution varied with depth and was dominated by Pelagibacterales and Rhodobacterales related Alphaproteobacteria in surface and intermediate layers, while Deltaproteobacteria increased in deep layers (up to 30%, Fig. S2). The relative abundance of Gammaproteobacteria was similar between water layers and dominated by members of Oceanospirillales. SAR11 clade related OTUs represented up to 60% of the total relative OTUs abundance (Fig.S2) and were dominant at all stations down to 300 m, regardless of the bloom regimes. Pelagibacterales or SAR11 clade, are the most abundant bacteria in marine ecosystem (Morris *et al.*, 2002) and have been shown to represent more than 25% of the total community during non-bloom conditions in the SO (West *et al.*, 2008; Giebel *et al.*, 2009)

The high contribution of the SAR11 clade to bulk bacterial abundance in surface waters was further observed by CARD-FISH (Table 1). The contribution of SAR11 cells to total bacterial abundance varied between 44.3±4% at Station R-2 HNLC waters and 33±10% at the three Fe-fertilized sites A3.2, F-L and E-5, in the surface and intermediate layers. In deep winter water layers, at 300m depth, the abundance of SAR11 cells strongly decreased (5 to 20% of contribution to total bacterial abundance). The success of SAR11 clade as the most abundant marine bacterial group, suggest that they play a major contribution to organic matter fluxes. Here, micro-autoradiography combined with CARD-FISH, showed that across the 300 m water column, the contribution of SAR11 cells to bulk bacterial abundance was correlated to its contribution to leucine incorporation (close to the 1:1 line, Fig.2A), indicating an overall high activity of SAR11 and important contribution to carbon cycling across the studied region at this time of the year. Similar ranges of SAR11 contribution to leucine incorporation were reported later in the season in HNLC waters (Obernosterer *et al.*, 2011). High activity has also been observed in other oceanic regions, such as the North Atlantic and Mediterranean Sea where SAR11 contributed to 30-50% of total leucine incorporation (Malmstrom *et al.*, 2004; Malmstrom *et al.*, 2005; Laghdass *et al.*, 2012).

**Figure 2:**
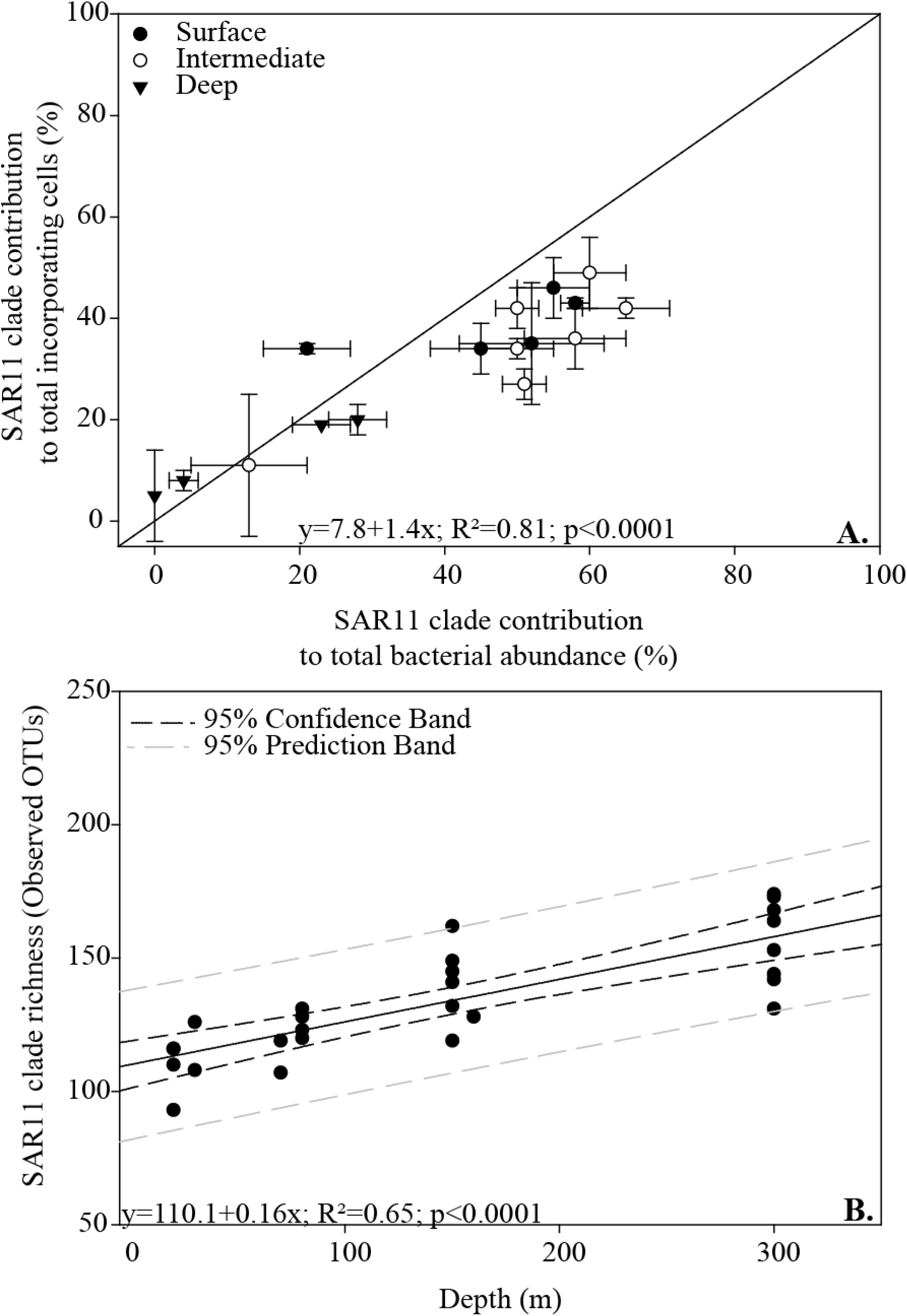
Contribution of SAR11 clade to total ^3^H-leucine incorporating cells versus contribution of SAR11 clade to total bacterial abundance. The solid line indicates a 1:1 relationship (A). Correlation between SAR11 clade richness (based on SAR11 clade related observed OTUs) and depth (B).

**Table 1:** SAR11 clade contribution to bacterial abundance and bacterial leucine incorporation.

| Station | Depth<br>(m) | BA<sup>1</sup><br>(x10<sup>5</sup> cells ml<sup>-1</sup>) | % cells of<br>SAR11<sup>2</sup> to<br>total BA | SAR11<br>abundance<br>(x 10<sup>4</sup> cells ml<sup>-1</sup>) | % of active<br>SAR11<sup>3</sup> cells | % of active<br>SAR11 cells to<br>total active<br>cells | % relative<br>abundance of<br>SAR11 OTUs |
| --- | --- | --- | --- | --- | --- | --- | --- |
| R-2 | 20 | 2.30 | 49 ± 7 | 11.2 | 51 ± 9 | 60 ± 5 | 47 |
| R-2 | 60 | 2.95 | 42 ± 4 | 12.4 | 38 ± 5 | 50 ± 3 | n.a. |
| R-2 | 150 | 2.87 | 42 ± 2 | 12.1 | 42 ± 3 | 65 ± 6 | 14 |
| R-2 | 300 | 1.30 | 19 ± 0 | 2.47 | 15 ± 3 | 23 ± 4 | 13 |
| A3-2 | 20 | 2.70 | 35 ± 12 | 9.45 | 12 ± 3 | 52 ± 10 | 36 |
| A3-2 | 80 | 3.53 | 43 ± 1 | 15.2 | 84 ± 3 | 58 ± 2 | 39 |
| A3-2 | 160 | 3.47 | 34 ± 1 | 11.2 | 58 ± 9 | 21 ± 6 | 41 |
| A3-2 | 300 | 1.90 | 20 ± 3 | 3.80 | 30 ± 6 | 28 ± 4 | 24 |
| F-L | 20 | 6.06 | 46 ± 6 | 27.9 | 37 ± 8 | 55 ± 5 | 31 |
| F-L | 70 | 6.48 | 34 ± 2 | 22.0 | 24 ± 4 | 50 ± 5 | 41 |
| F-L | 150 | 2.24 | 11 ± 14 | 2.46 | 3 ± 3 | 13 ± 8 | 34 |
| F-L | 300 | 1.81 | 5 ± 9 | 0.91 | 0 ± 1 | 0 | 19 |
| E-5 | 20 | 4.60 | 34 ± 5 | 15.6 | 60 ± 9 | 45 ± 7 | n.a. |
| E-5 | 80 | 4.56 | 36 ± 6 | 16.4 | 11 ± 4 | 58 ± 7 | 11 |
| E-5 | 150 | 3.57 | 27 ± 3 | 9.64 | 25 ± 7 | 51 ± 3 | 42 |
| E-5 | 300 | 2.20 | 8 ± 2 | 1.76 | 1 ± 1 | 4 ± 2 | 22 |

<sup>1</sup>BA: Bacterial abundance<sup>2</sup>SAR11 clade probes hybridized cells<sup>3</sup>Micro-autoradiography leucine incorporation positive cells

The high abundance and contribution to activity of Pelagibacterales at early bloom stages contrast with observations during the peak and decline of the phytoplankton bloom above the Kerguelen plateau (West *et al.*, 2008; Obernosterer *et al.*, 2011). Rhodobacterales (Alphaproteobacteria), Alteromonadales SAR92 (Gammaproteobacteria) and a member of Bacteroidetes (Agg58) were abundant and active members of the bacterial assemblage in late summer. These observations are in line with the idea that SAR11 is less competitive under high productivity conditions such as phytoplankton blooms (Malmstrom *et al.*, 2005; Tripp *et al.*, 2008). During the same cruise, SAR11 was experimentally shown to be rapidly outcompeted by other taxa in the presence of diatom-derived DOM (Landa *et al.*, 2018). The contribution of SAR11 cells to total abundance increased from 6 to 29% during the decline of the Kerguelen bloom (Obernosterer *et al.*, 2011). This suggests that while outcompeted during the peak of the bloom, SAR11 has the capacity to rapidly recover as the bloom ends, where abundances were similar to values observed early in the bloom (this study). The present and previous studies (Giebel *et al.*, 2009) highlight SAR11 to be major community members in non-productive Southern Ocean waters. Pelagibacterales are specialized in the utilization of labile low molecular weight DOM and outcompeted by other taxa for high molecular weight DOM utilization (Malmstrom *et al.*, 2005). This was also observed in our study region. While SAR11 cells were major contributors to leucine incorporation, they were less adapted at taking up more complex compounds such as chitin (Fourquez *et al.*, 2016). The success of SAR11 in these Fe-limited waters could further be due to strategies related to uptake, storage and utilization of this micronutrient (Debeljak *et al.*, 2019; Beier *et al.*, 2015).

The SAR11 lineage is a monophyletic group that contains several phylogenetic subclades, which have been shown to respond to environmental forcing and spatio-temperal variations in different oceanic regions (e.g. Brown *et al.*, 2012; Vergin *et al.*, 2013; Salter *et al.*, 2014). These subclades may represent niche specific ecotypes with different traits and metabolic needs (Grote *et al.*, 2012) which may in turn affect the water ecology and biogeochemistry. However, despite the success of SAR11 in the Southern Ocean waters, little is known about its microdiversity. In the present study, OTUs related to the SAR11 clade represented 40±15% of all bacterial OTUs across stations and depths layers (Fig.S2) and richness of SAR11 OTUs increased with depth (Fig.2B). SAR11 subclades distribution over the studied area and dynamics during the onset of the spring bloom, was further resolved to phylogenetic subclades partitioning (Fig. 3). Overall, 47 OTUs were closely related to Pelagibacterales at 99% identity (covering 64310 reads). The phylogenetic relationship between these OTUs and previously identified SAR11 subclasses (e.g. Field *et al.*, 1997; Carlson *et al.*, 2009; Vergin *et al.*, 2013) showed that OTUs observed in this study separated between six different known subclades. 80±6% of the reads belonged to subclade Ia, among which one OTU (KEOPS-6089) was especially abundant and represented 15% of the total relative abundance of all bacterial OTUs across all samples (Fig. 3). Subclade Ia is the most abundant and most studied SAR11 ecotype in the oceans (Giovannoni, 2017 and references therein). This subclade could be further separated in three subgroups (Fig. 3).

**Figure 3:**
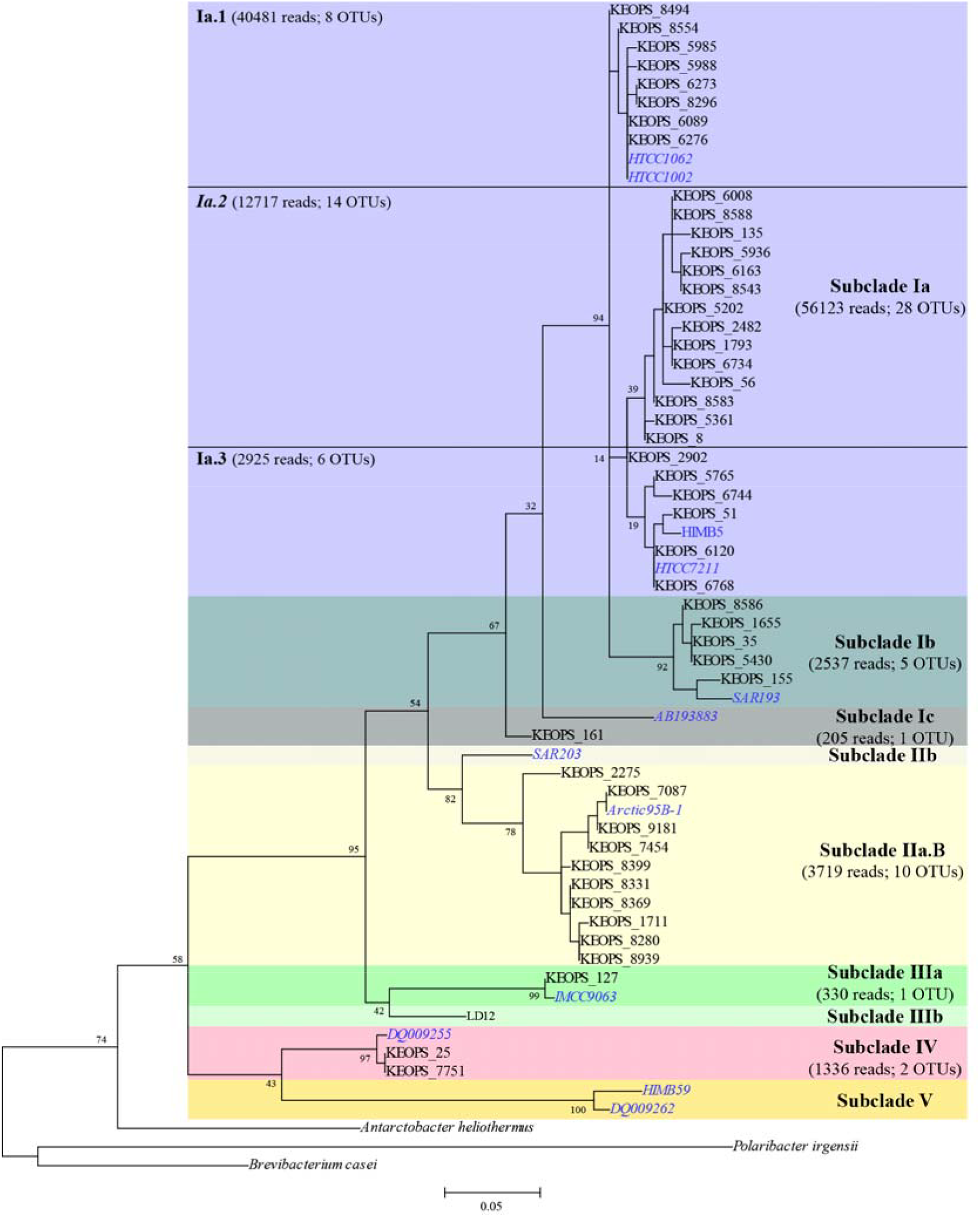
Phylogenetic relationships of SAR11 OTUs: Maximum likelyhood tree of OTUs closely related to SAR11 clade. Only SAR11 OTUs representing more than 0.1% of the total SAR11 reads are included. Reference sequences from previously published SAR11 subclades identifications are indicated in blue italic. Bootstrap values (n=1000) are indicated at nodes; scale bar represents changes per positions.

Because most of the SAR11 clades revealed minor variability in their distribution among sites, we describe in the following their depth distribution combining all stations. Specific SAR11 OTUs were more represented in different depth layers (Fig. 4). More specifically, subclades Ia, IIIa and IV were relatively more abundant in surface and intermediate layers, while subclade Ib was more abundant in the deep layers (Fig. 4.A). At the Atlantic time series site BATS, subclade Ib is usually found in surface water in late spring and early summer, while Ic is found in deeper water (Vergin *et al.*, 2013). Subclade Ib has also been reported in epi- and bathypelagic waters in the Red Sea where it blooms following water mixing (Jimenez-Infante *et al.*, 2017). Subgroup Ia.3 was more abundant in the deep layers, while Ia.1 and Ia.2 had higher relative abundances in surface and intermediate layers (Fig. 4.B). Ia.1 has been observed in colder surface coastal waters (Rappe *et al.*, 2002; Brown *et al.*, 2012; Grote *et al.*, 2012), while Ia.3 has been reported in surface gyre and tropical waters (Brown *et al.*, 2012; Grote *et al.*, 2012). Nevertheless, Ia.3 has also been observed in deep water of the Red Sea (Ngugi and Stingl, 2012). Differences in the SAR11 subclade distributions were most pronounced between the surface/intermediate and deep layers, an observation that could possibly be linked to the different water masses that are the Antarctic and Polar Front surface and winter waters. The different distribution of these subclades in the present study (such as Ib, Ic and Ia3) may be due to the water masses origin and formation specific to the Southern Ocean.

**Figure 4:**
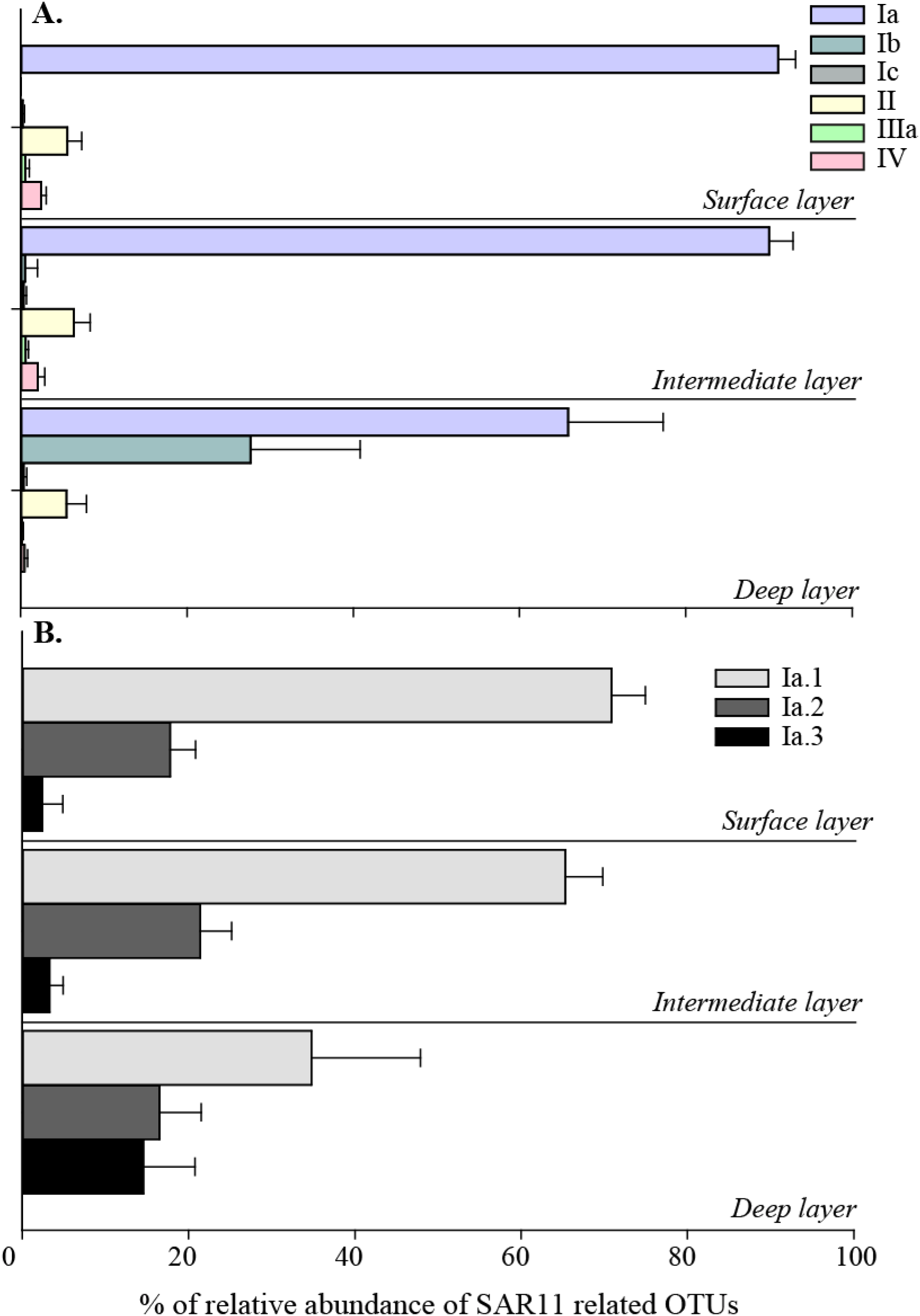
Layer distribution of SAR11 subclades (A) and subclades Ia (B): % relative abundance of SAR11 related OTUs representing more than 0.1 % of all SAR11 OTUs. Average of SAR11 relative abundance across all stations. Error bars represent standard deviation between stations.

The microdiversity of SAR11 subclades appeared more dynamic in surface waters (20m), similar to the shift in surface bacterial community composition following the bloom progression (Landa *et al.*, 2016). Despite the separate clustering of surface and intermediate waters based on environmental conditions, only few of the different subclades revealed specific patterns between these water layers (Fig. 4) and bloom stages (Fig S3, S4). For instance subclade IV was most abundant in more advanced bloom stage stations (A3.2, E4W and FL), while subclade Ic and IIa seemed most abundant in the non-bloom stations (R-2, HNLC station and E3, Fig. S3). In tropical regions subclade IV is associated to summer deep chlorophyll maximum waters, while IIa is found in surface winter and spring waters (Vergin *et al.*, 2013). The bi-polar distribution of subclade IIa in surface waters has been previously reported (Kraemer *et al.*, 2019), its adaptation to cold waters may explain its relatively stable distribution in less productive stations across the water column in the study area. Subclades Ia and Ib did not seem to respond to bloom stages (Fig. S3). Ia.2 appeared to be more abundant in the HNLC waters, this previously uncharacterized subgroup may be locally adapted to less productive and Fe-limited waters of the SO. The spatio-temporal partitioning of some of the SAR11 subclades revealed in this study suggest a niche specificity and periodic selection of the different SAR11 ecotypes in the SO, which may in turn impact the role of SAR11 in biogeochemical cycles throughout the bloom progression.

In conclusion, Pelagibacterales appear highly adapted to the cold HNLC waters of the Southern Ocean where they contribute substantially to bacterial biomass production and probably other microbially-mediated fluxes. Future studies should focus on ITS phylotypes and isolations of different SAR11 representative in order to better constrain the microdiversity, as well as the evolution of locally adapted strains and their ecological function in the Southern Ocean.

## Experimental procedures

### Study site and sampling

The sampling was conducted during the KEOPS2 (Kerguelen Ocean and Plateau Compared Study 2) cruise from October to November 2011 on board the R/V Marion Dufresne in the Kerguelen region (FigS1 and see Fig. 1 in Landa *et al.*, 2016). A total of seven stations (A3.2, E stations and F-L) were sampled in the naturally iron-fertilized regions east of the Kerguelen Islands and a reference station (R-2) was sampled in high nutrient low chlorophyll waters (HNLC) located west of the islands. For each station four depths were sampled according to CTD profiles.

### Bacterial community composition

A total of 31 samples from eight different stations were analyzed for bacterial community composition. Filtration, extraction, sequencing of the V1-V3 16S rDNA and denoising of the sequences are described in Landa *et al.* (2016). Clean reads were subsequently processed using the Quantitative Insight Into Microbial Ecology pipeline (QIIME v1.7; (Caporaso *et al.*, 2010b)). Reads were clustered into (OTUs) at 99% pairwise identity using Uclust and representative sequences from each bacterial OUT were aligned to Greengenes reference alignment using PyNAST (Caporaso *et al.*, 2010a). All singletons and operational taxonomic units (OTUs) present in only one sample were removed. Taxonomy assignments were made using the Ribosomal Database Project (RDP) classifier (Wang *et al.*, 2007) against the database Greengene 13_8 (McDonald *et al.*, 2012) and SILVA 128 (Quast *et al.*, 2013). SAR11 maximum likelyhood tree was computed with Mega7 (Tamura *et al.*, 2013). The surface sample from station E5 was removed from the dataset due to unsatisfactory quality results. The data were deposited in the Sequence Read Archive (SRA) database under accession number SRP041580. Alpha and beta-diversity were estimated using QIIME for all samples after randomized subsampling to 3166 reads for the total diversity and to 500 reads for SAR11 clade diversity, details are described in Supplementary material.

### Micro-CARD-FISH

Leucine uptake by SAR11 cells was studied through Micro-CARD-FISH. The details of the method are described in (Fourquez *et al.*, 2016). Briefly, 10 mL of seawater samples were incubated with radiolabeled leucine for 6-8h, fixed with paraformaldehyde and filtered onto 0.2 µm polycarbonate filters. The abundance of SAR11 was determined with catalyzed reporter deposition fluorescence in situ hybridization (CARD-FISH) with the probes SAR11-152R, SAR11-441R, SAR11-542R and SAR11-732R (Morris *et al.*, 2002). The micro-autoradiography development with photographic emulsion was exposed for 2-3d. After development the proportion of substrate active SAR11 were determined as the proportion of probe positive cell with silver grains.

## Acknowledgments

We thank S. Blain, the PI of the KEOPS2 project, for providing us the opportunity to participate to this cruise, the chief scientist B. Quéguiner, the captain Bernard Lassiette and the crew of the R/V Marion Dufresne for their enthusiasm and help aboard. This work was supported by the French Research program of the INSU-CNRS LEFE−CYBER (Les enveloppes fluides et l’environnement –Cycles biogéochimiques, environnement et ressources), the French ANR (Agence Nationale de la Recherche, SIMI-6 program), the French CNES (Centre National d’Etudes Spatiales) and the French Polar Institute IPEV (Institut Polaire Paul−Emile Victor). JD was supported by the Marie Curie Actions-International Outgoing Fellowship (PIOF-GA-2013-629378).

