## Supplementary material for "High contribution of *Pelagibacterales* to bacterial community composition and activity in spring blooms off Kerguelen Island (Southern Ocean)"

**Supplementary experimental procedures**

After quality trimming, a total of 201357 reads distributed over 2973 OTUs (Operational taxonomic units, at 99% identity), were retrieved from 31 samples collected in the upper 300 m of the water column with an average of 6292 reads per samples. Alpha and beta-diversity indices were estimated for all samples after randomized subsampling to 3166 reads. Alpha diversity analyses (Richness: observed OTUs and diversity: Shannon index) were run in QIIME. Beta-diversity and link to environmental parameters were investigated with Primer V.6 software package (Clarke and Warwick, 2001). Euclidan distances between the samples environmental parameters (main environmental variables see table S2) were analyzed by non-metric multidimensional scaling (nMDS) of log X+1 normalized datasets. The similarity between bacterial community compositions in the samples was analyzed with nMDS of Bray-curtis dissimilarities on log X+1 transformed dataset. The similarity profile (SIMPROF) permutation tests were conducted to establish the significance of dendrogram branches resulting from the cluster analyses. Correlation between resemblance matrices of bacterial community and environmental parameter was tested with Spearman rank correlation.

**Supplementary Table and Figures**

**Supplementary Table S1:** Depth layer classification, environmental and bacterial parameters for the sampled stations. Bacterial taxonomic richness is expressed as number of OTUs, bacterial diversity is calculated with Shannon Index. (n.a.: not available, HLNC: High nutrients low chlorophyll station). Mixed layer depths, based on a difference in sigma of 0.02 to the surface value, are 105 m at R-2, 38m at F-L, 61m at E4-W, 153m at A3.2, 72m at E1, 38m at E3, 74 m at E4E, 46m at E5. Layers are determined based on the environmental data clustering in Figure 1A.

| **Station group** | **Station** | **Depth (m)** | **layer** | **Chl. *a* (µg L^-1^)*** | **Bacterial abundance (x10^5^ cells mL^-1^)*** | **Bacterial production (ng C L^-1^ h^-1^)*** | | **Bacterial richness** | | | **Bacterial diversity** |
| --- | --- | --- | --- | --- | --- | --- | --- | --- | --- | --- | --- |
| HNLC | **R-2** | 20 | Intermediate | 0.32 | 2.29 | | 2.08 | | 423 | 6.27 | |
|  | **R-2** | 60 | Intermediate | 0.27 | 2.95 | | 2.77 | | 242 | 5.39 | |
|  | **R-2** | 150 | Intermediate | 0.07 | 2.87 | | 0.88 | | 344 | 6.31 | |
|  | **R-2** | 300 | Deep | - | 1.30 | | 0.63 | | 493 | 6.92 | |
| Polar front plume | **F-L** | 20 | Surface | 5.12 | 6.06 | | 64.6 | | 370 | 6.21 | |
|  | **F-L** | 70 | Intermediate | 0.34 | 6.48 | | 6.82 | | 435 | 6.38 | |
|  | **F-L** | 150 | Intermediate | 0.04 | 2.24 | | 1.49 | | 636 | 7.41 | |
|  | **F-L** | 300 | Deep | - | 1.81 | | 0.23 | | 536 | 7.13 | |
| Kerguelen  Plateau | **E4W** | 30 | Surface | 1.40 | 6.04 | | 30.16 | | 427 | 5.93 | |
|  | **E4W** | 80 | Surface | 1.22 | 5.96 | | 24.83 | | 420 | 6.18 | |
|  | **E4W** | 150 | Intermediate | 0.22 | 3.15 | | 4.34 | | 502 | 6.74 | |
|  | **E4W** | 300 | Deep | - | 1.76 | | 0.31 | | 606 | 7.44 | |
|  | **A3.2** | 20 | Surface | 1.65 | 2.70 | | 20.15 | | 433 | 6.43 | |
|  | **A3.2** | 80 | Surface | 2.12 | 3.53 | | 20.63 | | 402 | 6.30 | |
|  | **A3.2** | 160 | Surface | 2.30 | 3.47 | | 22.43 | | 449 | 6.57 | |
|  | **A3.2** | 300 | Deep | - | 1.90 | | 1.09 | | 620 | 7.52 | |
| Recirculation  feature | **E1** | 20 | Surface | 1.00 | 4.33 | | 15.98 | | 433 | 6.45 | |
|  | **E1** | 80 | Intermediate | 0.85 | 4.26 | | 14.73 | | 484 | 6.52 | |
|  | **E1** | 150 | Intermediate | 0.60 | 3.83 | | 9.78 | | 461 | 6.29 | |
|  | **E1** | 300 | Deep | - | 1.34 | | 0.25 | | 610 | 7.60 | |
|  | **E3** | 20 | Surface | 0.69 | 5.06 | | 23.65 | | 309 | 5.44 | |
|  | **E3** | 70 | Intermediate | 0.42 | 4.93 | | 16.6 | | 448 | 6.52 | |
|  | **E3** | 150 | Intermediate | 0.50 | 4.18 | | 9.02 | | 505 | 6.88 | |
|  | **E3** | 300 | Deep | - | 1.81 | | 0.48 | | 645 | 7.63 | |
|  | **E4E** | 30 | Surface | 1.09 | 5.63 | | 39.65 | | 413 | 6.32 | |
|  | **E4E** | 80 | Intermediate | 0.39 | 5.28 | | 10.97 | | 484 | 6.96 | |
|  | **E4E** | 150 | Intermediate | 0.19 | 3.18 | | 5.06 | | 542 | 6.79 | |
|  | **E4E** | 300 | Deep | - | 1.67 | | 0.17 | | 721 | 8.01 | |
|  | **E5** | 20 | Surface | 1.21 | 4.60 | | 28.27 | | n.a. | n.a. | |
|  | **E5** | 80 | Intermediate | 0.92 | 4.56 | | 26.43 | | 326 | 5.52 | |
|  | **E5** | 150 | Intermediate | 0.20 | 3.57 | | 3.3 | | 561 | 7.03 | |
|  | **E5** | 300 | Deep | - | 2.20 | | 0.15 | | 754 | 8.00 | |

*data from Christaki et al. 2014

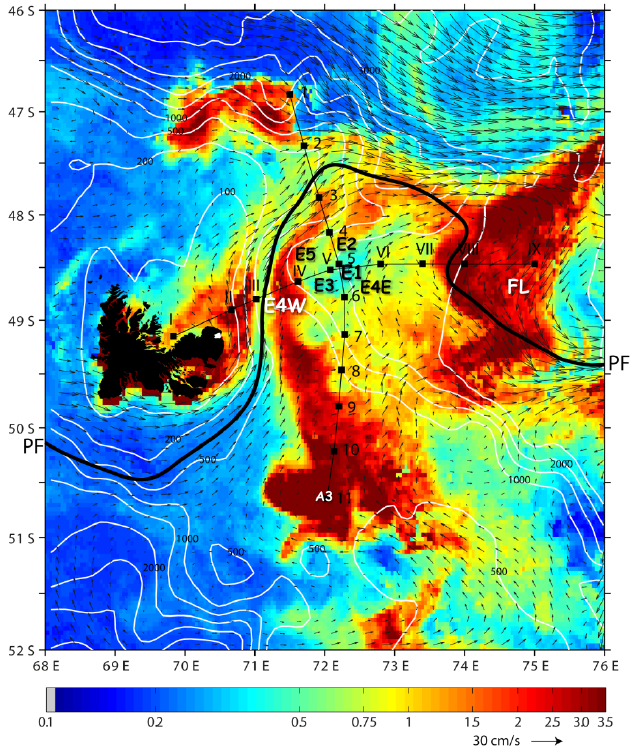

**Supplementary Figure S1:**  KEOPS2 study area from October to November 2011. Chl a (colour scale), surface velocity fields (arrows), the polar front (PF, black line), and the position of the different stations: The Chl *a* rich stations: A3, on the Kerguelen plateau; F-L and E-4W north and south of the polar front; and “E” stations sampled in a quasi-Lagrangian manner (E-1, E-2, E-3, E-4E, and E-5) within a complex meander south of the polar front. The reference HNLC station (R-2) is not shown as it is out of the area of the map (66.692743 E longitude, 50.38954 N latitude). Map is courtesy of Y. Park and colleagues. To note: the chlorophyll content represented on the map corresponds to the last week of the KEOPS2 cruise.

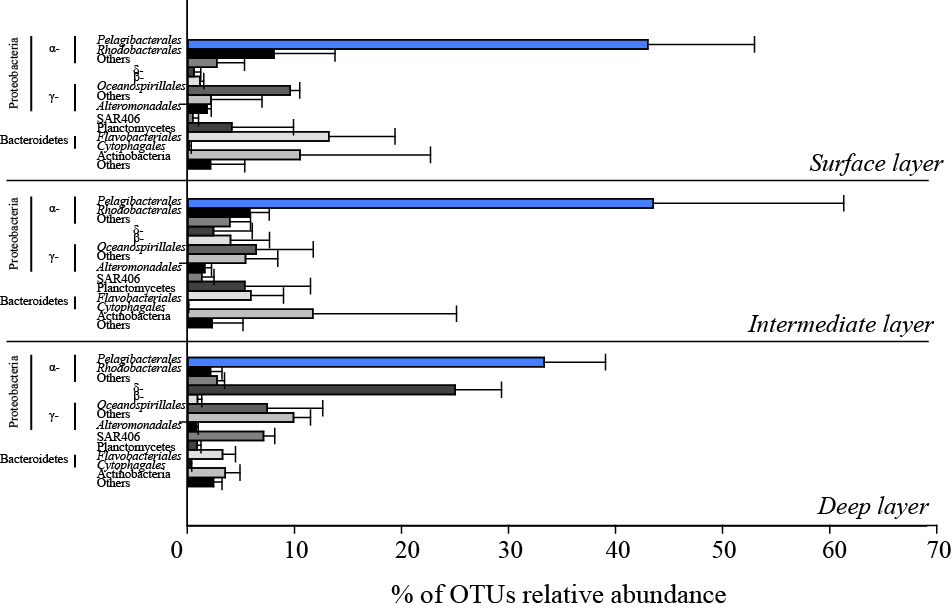

**Supplementary Figure S2:** Relative abundance (%) of the main taxonomic groups over the three depth layers

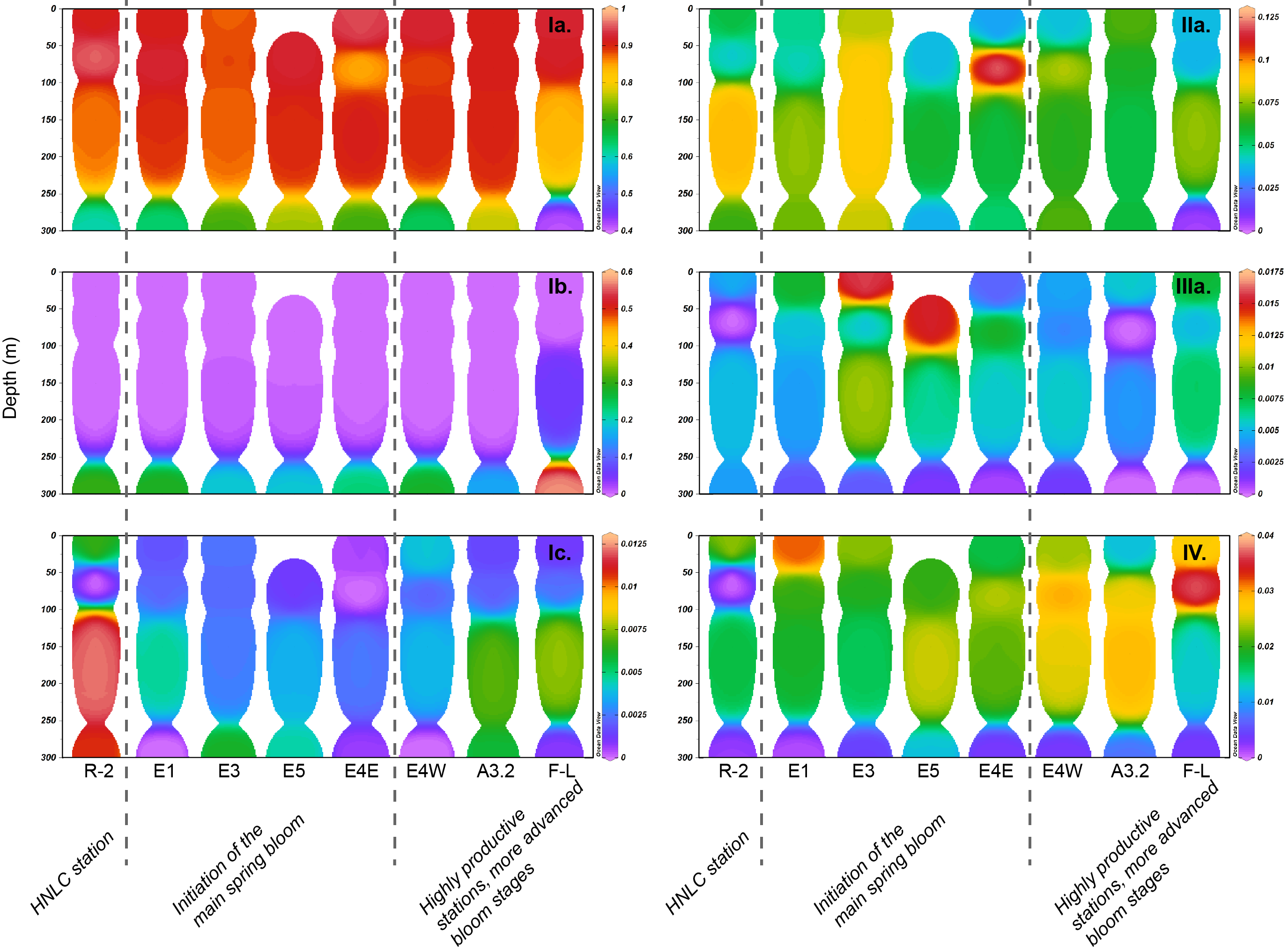

**Supplementary Figure S3:** Vertical distribution of SAR11 subclades under different bloom conditions (integrated weight average of SAR11 relative abundance to total community, note different z scales).

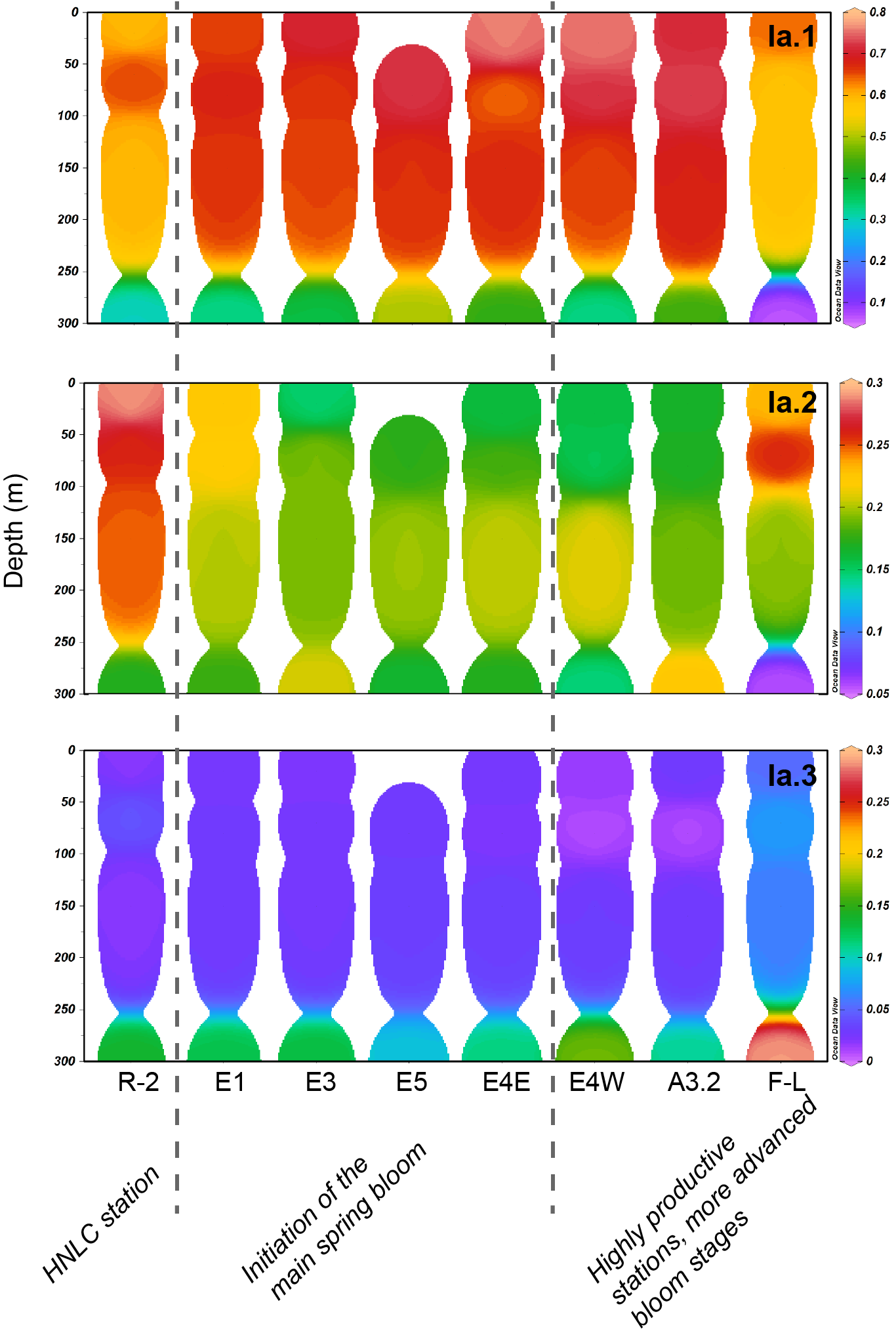

**Supplementary Figure S4:** Vertical distribution of SAR11 subclades Ia. under different bloom conditions (integrated weight average of SAR11 relative abundance to total community, note different z scales).
